# Estimating serotype-specific efficacy of pneumococcal conjugate vaccines using hierarchical models

**DOI:** 10.1101/633909

**Authors:** Joshua L Warren, Daniel M. Weinberger

## Abstract

Pneumococcal conjugate vaccines (PCVs) target 10 or 13 specific serotypes. To evaluate vaccine efficacy for these products, the vaccine-targeted serotypes are typically aggregated into a single group to estimate an overall effect. However, it is often desirable to evaluate variations in effects for different serotypes. These serotype-specific estimates are often based on small numbers, resulting in a high degree of uncertainty and instability in the individual estimates. A better approach is to use a Bayesian hierarchical statistical model, which estimates an overall effectiveness of the vaccine across all vaccine-targeted serotypes but also allows the effect to vary by serotype. We re-analyzed published data from a large randomized controlled trial on the efficacy of PCV13 against non-bacteremic community-acquired pneumonia caused by vaccine-targeted serotype. This model provides a potential framework for obtaining more credible and stable estimates of serotype-specific vaccine efficacy and effectiveness.

## Introduction

Pneumococcus is a diverse bacterial pathogen, with more than 90 identified serotypes. Pneumococcal conjugate vaccines (PCVs) protect against a subset of these serotypes, with currently-available vaccines targeting 10 or 13 serotypes, and next-generation vaccines targeting 15 or 20 serotypes. To evaluate vaccine efficacy or effectiveness for these products, the vaccine-targeted serotypes are typically aggregated into a single group to estimate an overall effect [1, 2]. However, it is often desirable to evaluate variations in effects for different serotypes [3, 4]. The problem is that studies are not typically powered to obtain reliable stratified estimates. As a result, the serotype-specific estimates are often based on small numbers, resulting in a high degree of uncertainty and instability in the individual estimates. This can lead to incorrect interpretations, such as concluding that the effect the vaccine against individual serotypes is too large or too small.

Analyses of vaccine effects typically follow one of two approaches: they pool all vaccine serotypes together, assuming that the vaccine serotypes are interchangeable; or they estimate effectiveness separately for each serotype individually but ignore the estimates for the other serotypes. A compromise approach is to use a hierarchical statistical model, which estimates an overall effectiveness of the vaccine across all vaccine-targeted serotypes but also allows the effect to vary by serotype. With this approach, the estimate of effectiveness obtained for each serotype is somewhere between the overall average effect and the effect that would be estimated by analyzing each serotype individually. For serotypes with more robust data, the estimate will fall closer to the estimate that would be obtained by analyzing each serotype individually, while for serotypes with few cases, the estimate will fall closer to the overall estimate.

In this study, we re-analyzed serotype-specific vaccine efficacy data from the CAPITA trial, which was a randomized controlled trial that evaluated the effects of PCV against pneumococcal pneumonia [5]. This hierarchical model provides a potential framework for obtaining more credible and stable estimates of serotype-specific vaccine efficacy and effectiveness.

## Methods

### Data

The data were extracted from the CAPITA randomized controlled trial published by Bonten, et al. [5]. Adults were randomized to receive either the 13-valenet pneumococcal conjugate vaccine (PCV13) or a placebo. Our analyses focus on the data presented in Table S11: number of cases of first episode non-bacteremic community-acquired pneumonia (CAP) caused by a serotype targeted by PCV13, according to the per protocol analysis. The denominator was 42,240 in the vaccinees and 42,256 in the controls.

### Overall estimate of efficacy

The first analysis is a multinomial logistic regression where the outcome is the number of cases of each of the 13 vaccine-targeted serotypes and the number of people who did not develop CAP (‘non-cases’) in the vacinees and in the controls. We first assumed that the vaccine efficacy was the same for all targeted serotypes such that

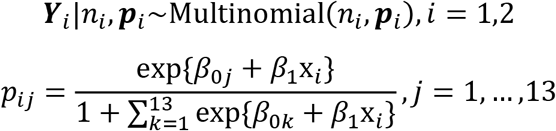

where ***Y***_*i*_ is a vector that contains the number of cases due to each serotype (*Y*_*i*1_, …, *Y*_*i*13_) and the number of non-cases (*Y*_*i*14_) for treatment group *i* (*i*=1: placebo, *i*=2: PCV13), *n*_*i*_ is the number of study participants in treatment group *i* where the sum of the counts across all categories of ***Y***_*i*_ is equal to *n*_*i*_, and x_*i*_ is an indicator for vaccine status (x_1_=0: placebo, x_2_=1: PCV13). The probabilities that control how many people in treatment group *i* are detected in each category is defined by the ***p***_*i*_ vector, which has a unique entry for each vaccine-targeted serotype and non-case category, *p*_*ij*_, *j*=1,…,14. These probabilities sum to one for each treatment group so that the probability of not developing disease is given as 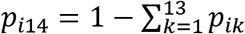. Vaccine efficacy serotype *j* (*j*=1,…,13) is defined as 100(1 − *p*_2*j*_/*p*_1*j*_) and the overall efficacy is defined as 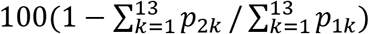. In this case, all estimated effects are the same since the vaccine efficacy parameter (*β*_1_) is shared across all serotypes. We fit the model in the Bayesian setting while specifying weakly informative, independent normally distributed prior distributions for the regression parameters (mean 0, standard deviation (sd) 100).

### Serotype-specific estimates of efficacy

Building on the model above, we now allow the vaccine effect to vary by serotype such that

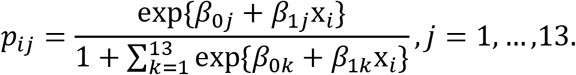

We evaluated two versions of this model: one where the serotype-specific intercepts (*β*_0*j*_) and slopes (*β*_1*j*_) were estimated independently for each serotype and one where they were estimated hierarchically. For the *non-hierarchical* version, each *β*_0*j*_ and *β*_1*j*_ were assigned independent, weakly informative normal priors (mean 0, sd 100). For the hierarchical model, these regression parameters were once again assigned independent, normally distributed prior distributions. However, the means and variances of these prior distributions were treated as unknown parameters; 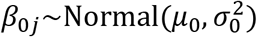 and 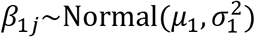, *j*=1,…13. These hyper-parameters were then assigned weakly informative priors to allow the data to determine the appropriate amount of shrinkage between the two extreme scenarios considered previously (shared vaccine effect, varying vaccine effect but non-hierarchical) such that *μ*_0_, *μ*_1_~Normal(0,100^2^) and *σ*_0_, *σ*_1_~Uniform(0,100). Vaccine efficacy was estimated as previously detailed.

All model fitting was performed using JAGS with the rjags package [6], using three separate Markov chains with a burn-in period of 10,000 iterations in each chain. Posterior inference was based on a total of 300,000 samples (100,000 from each chain) collected post-convergence, and vaccine efficacy was summarized using posterior medians as point estimates and 95% highest posterior density intervals to quantify uncertainty. Forest plots were generated with the rmeta package [7]. Analyses were performed in RStudio with the R statistical software [8]. All code and data are available on a public github repository: https://github.com/weinbergerlab/capita-hierarchical.

## Results

### Overall estimate of efficacy

There were 61 cases of vaccine-type pneumococcal pneumonia in the control group and 33 cases in the vaccine group. Without taking serotype into account, the estimated efficacy was 44.9% (95% credible intervals (CrI): 20.6%, 66.8%). This is close to the efficacy estimate reported in the original study (45.0%, 95% confidence interval: 14.2%, 65.3%).

### Serotype-specific efficacy, estimated separately

Only three of the serotypes: 3, 7F, and 19A had enough cases to obtain a reasonable estimate of VE when analyzed independently (Figure 1). Even for these serotypes, the estimates were based on fewer than 15 cases per group. The estimates for the remaining 10 serotypes were highly unstable. Many of the serotypes had 0 or 1 cases in either the vaccine or control group, which led to estimation instability. For instance, the estimates for serotypes 14, 19F and 23F were 100% because there were 0 cases in the vaccine group and only 2 cases in the control group. Serotype 18C, with 4 cases each in the case and control group, had an estimated efficacy of −0.4%, with wide CrIs (−246%, 92%). And some serotypes had point estimates of vaccine efficacy that were strongly negative. For instance, serotype 6A had 4 cases in the vaccine group and 2 in the control group, for a VE of −118% (95% CrI: −1087%, 92%). No reasonable estimate could be obtained for serotype 9V (1 case in vaccine group, 0 in control group). The overall estimate of vaccine efficacy with this model was 46.1% (95% CrI: 21.2%, 67.2%).

**Figure 1.**
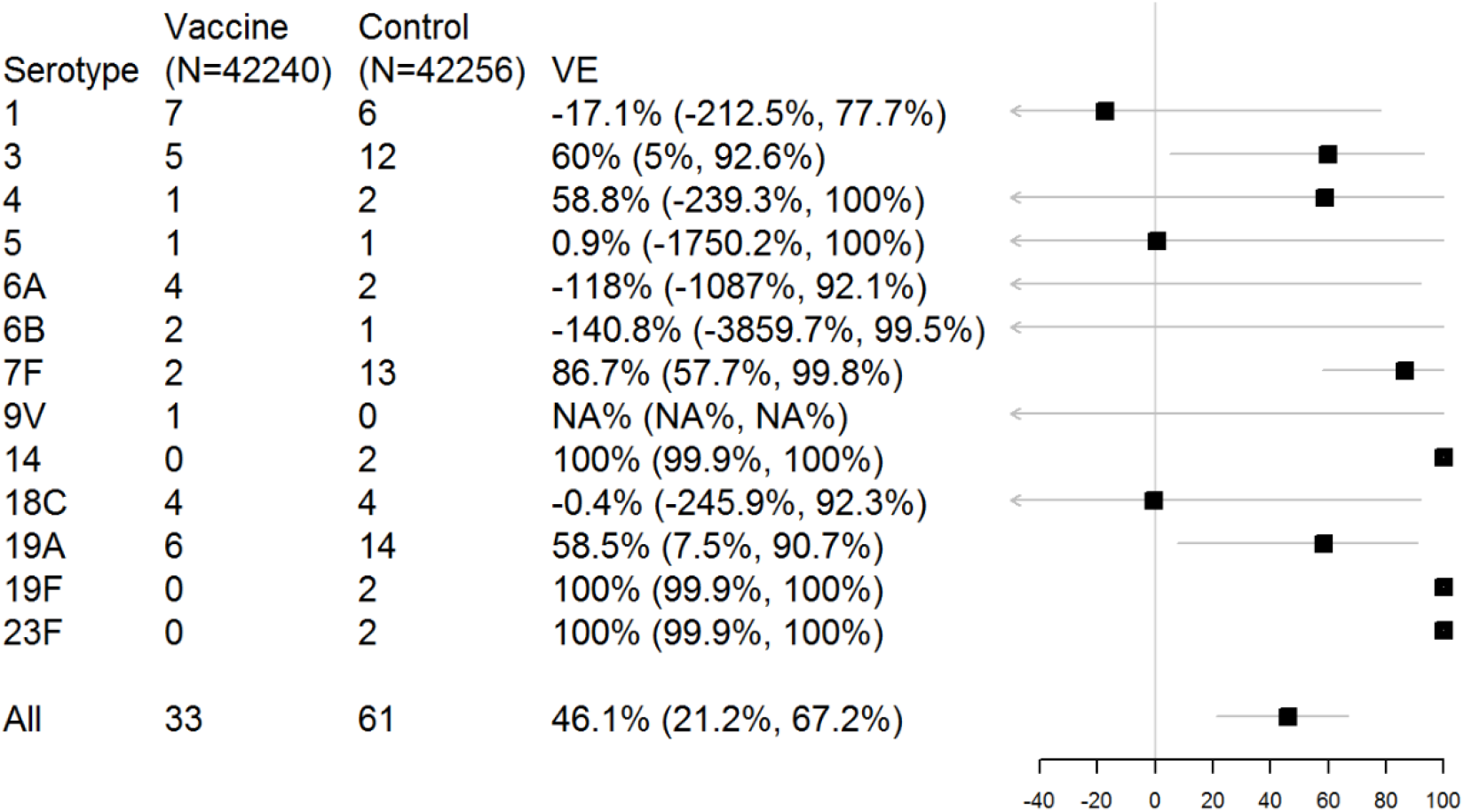
Estimates of serotype-specific vaccine efficacy from a non-hierarchical model. The plot shows the posterior median and 95% highest posterior density credible intervals.

### Serotype-specific efficacy, estimated hierarchically

Compared to the estimates shown in Figure 1, the serotype-specific effects were pulled towards the overall estimate (Figures 2 and 3), and serotypes with fewer observed cases were pulled more strongly towards the overall estimate (Figure 3). For serotypes where the observed data were consistent with the overall average, even if there were few counts, the CrIs were narrow and did not include 1 (Serotypes 3, 7F, 14, 19A, 19F, 23F; Figure 2). In contrast, for serotypes 1, 6A, 18C, more cases occurred among the vaccine recipients than the controls (based on few cases). For these serotypes, the point estimates of vaccine efficacy were pulled towards 0, with wider CrIs. The overall estimate of vaccine efficacy with this model was 46.0% (21.2%, 67.3%).

**Figure 2.**
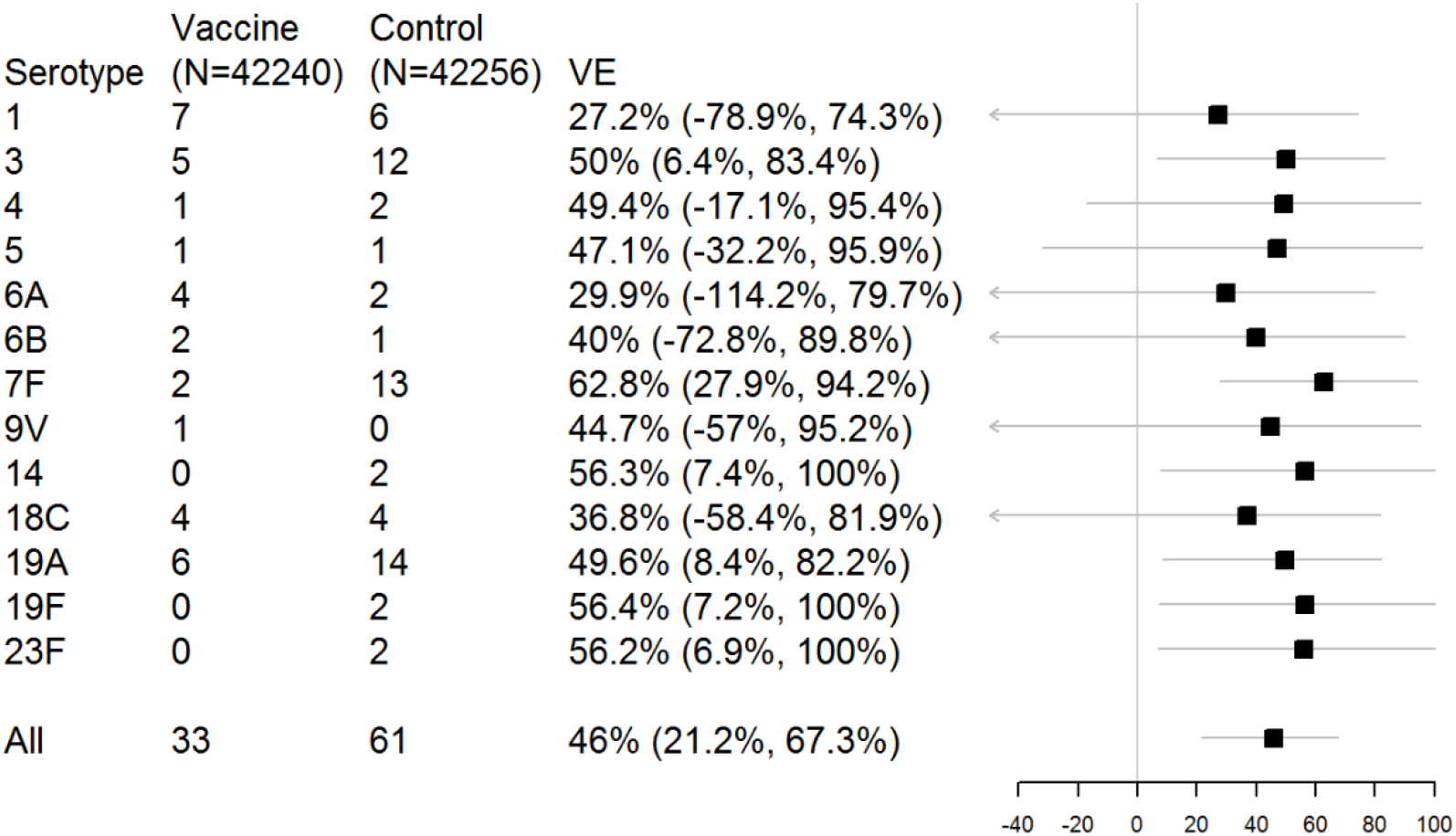
Estimates of serotype-specific vaccine efficacy from a hierarchical model. The plot shows the posterior median and 95% highest posterior density credible intervals.

**Figure 3.**
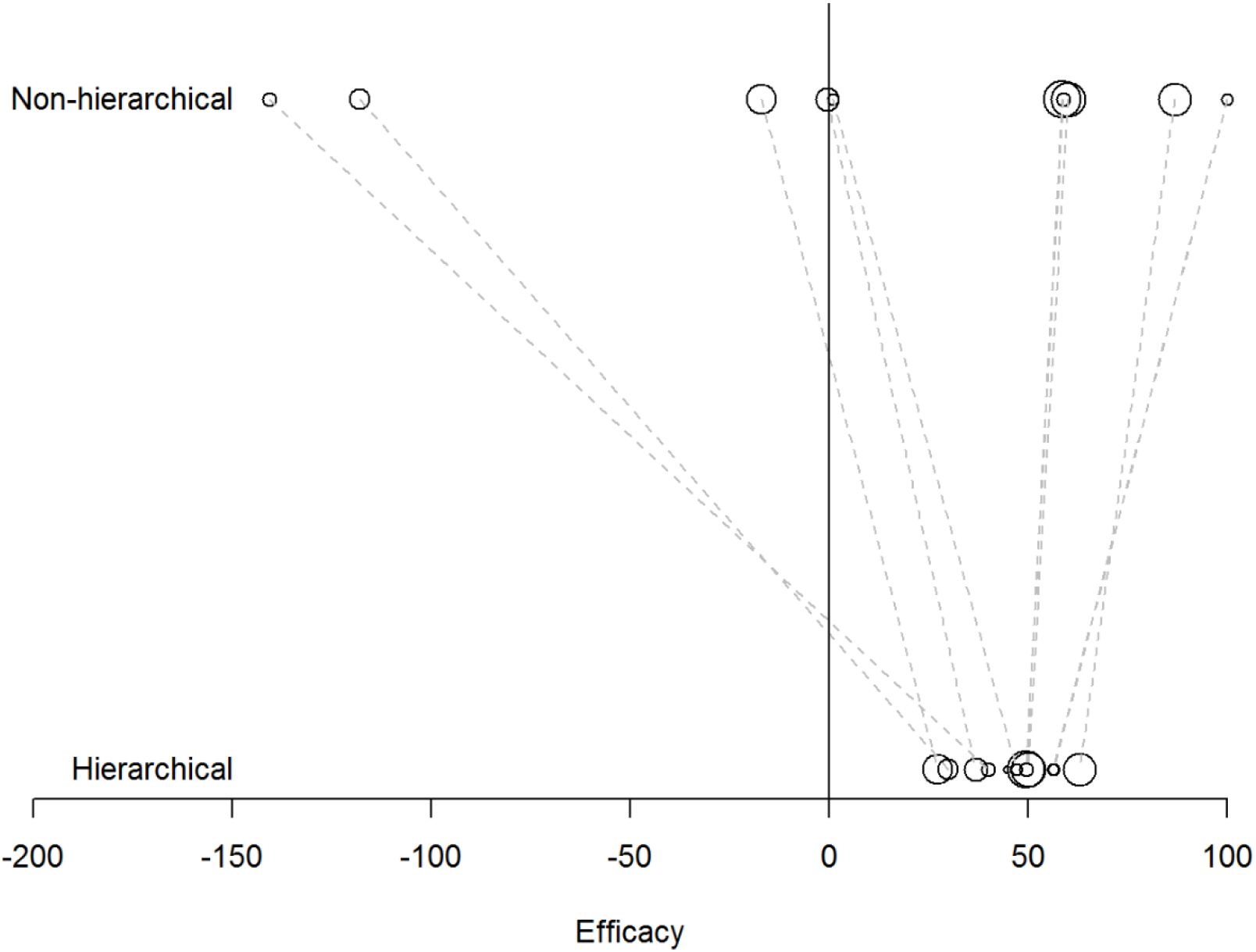
Comparison of the point estimates (posterior medians) of vaccine efficacy for each serotype from the non-hierarchical model and the hierarchical model. The area of the bubble is proportional to the number of cases of pneumonia due to that serotype in both study arms.

## Discussion

We present a framework to estimate serotype-specific vaccine effects. This hierarchical approach allows for more stable estimates of efficacy for each serotype while allowing the estimates to vary by serotype. As more data accrue from other trials or observational studies, the serotype-specific estimates could be updated. This approach could be used to evaluate any vaccine where there is expected to be variation in efficacy by strain.

The same general model structure that was used to analyze the data from an RCT in this study could be used to estimate serotype-specific effectiveness using observational data from a case-control study. Individual-level covariates (e.g., age) could be incorporated, and the associations could vary by serotype.

A strength and limitation of this type of hierarchical model is that the estimates for each serotype are pulled towards an overall mean. This shrinkage can be a desirable property and often reduces the overall mean-squared error of the estimators by introducing bias in order to achieve a reduction in variance. The impact of the shrinkage is typically greater for serotypes with fewer data.

In conclusion, we present an analytical framework for obtaining stable estimates of serotype-specific vaccine efficacy and effectiveness. This approach balances the need for serotype-specific estimates with the challenges of using sparse data that are subject to random variation. This approach has the potential to provide more credible estimates that can be compared between settings.

